# SARS-CoV-2 highly conserved s2m element dimerizes via a kissing complex and interacts with host miRNA-1307-3p

**DOI:** 10.1101/2020.12.29.424733

**Authors:** Joshua A. Imperatore, Caylee L. Cunningham, Kendy A. Pellegrene, Robert G. Brinson, John P. Marino, Jeffrey D. Evanseck, Mihaela Rita Mihailescu

## Abstract

The ongoing COVID-19 pandemic highlights the necessity for a more fundamental understanding of the coronavirus life cycle. The causative agent of the disease, SARS-CoV-2, is being studied extensively from a structural standpoint in order to gain insight into key molecular mechanisms required for its survival. Contained within the untranslated regions of the SARS-CoV-2 genome are various conserved stem-loop elements that are believed to function in RNA replication, viral protein translation, and discontinuous transcription. While the majority of these regions are variable in sequence, a 41-nucleotide s2m element within the 3’ UTR is highly conserved among coronaviruses and three other viral families. In this study, we demonstrate that the s2m element of SARS-CoV-2 dimerizes by forming an intermediate homodimeric kissing complex structure that is subsequently converted to a thermodynamically stable duplex conformation. This process is aided by the viral nucleocapsid protein, potentially indicating a role in mediating genome dimerization. Furthermore, we demonstrate that the s2m element interacts with multiple copies of host cellular miRNA-1307-3p. Taken together, our results highlight the potential significance of the dimer structures formed by the s2m element in key biological processes and implicate the motif as a possible therapeutic drug target for COVID-19 and other coronavirus-related diseases.

## INTRODUCTION

In late December 2019, a novel respiratory infection emerged in Wuhan, China and was consequently termed coronavirus disease 2019 (COVID-19) by the World Health Organization. The COVID-19 outbreak was declared a pandemic approximately 3 months later and has, to date, spread to over 200 countries with more than 71 million confirmed cases and over 1.6 million deaths (1). Symptoms of COVID-19 include those commonly seen in other respiratory diseases, such as fever, cough, and fatigue, though various neurological symptoms, including headache and dizziness, have also been reported (2). Severe acute respiratory syndrome coronavirus 2 (SARS-CoV-2), the virus responsible for COVID-19, is proposed to have originated by homologous recombination events between bat and pangolin coronaviruses, which most likely enabled its transmission to humans (3, 4). Though high rates of homologous recombination have been reported for coronaviruses, the mechanism by which this process occurs remains elusive (5).

SARS-CoV-2 is a member of the Nidovirales order, *Coronaviridae* family, and *Betacoronavirus* genus, lineage B (6). Similar viruses, such as SARS coronavirus (SARS-CoV) and Middle East respiratory syndrome (MERS) coronavirus, resulted in high fatality rates among infected individuals during the SARS and MERS outbreaks in 2002 and 2012, respectively (7). Members of the *Coronaviridae* family are enveloped, single-stranded RNA viruses with positive polarity and large genomes of approximately 30 kilobases (8, 9). These viruses contain similar genomic RNA (gRNA) composition, including two open reading frames (ORF1a and ORF1b), which encode for the RNA-dependent RNA polymerase (RdRp) and nonstructural proteins (10). Interestingly, members of the *Coronaviridae* family of viruses harbor the unique ability to produce “nested” subgenomic RNAs (sgRNA), which serve as the templates for translation of the virion’s structural spike (S), membrane (M), envelope (E), and nucleocapsid (N) proteins (10, 11). The latter of these proteins, N, forms ribonucleoprotein particles with the gRNA, termed the nucleocapsid complex, which is enveloped by a membrane consisting of S, E, and M proteins during the virion maturation process.

The genomes of *Coronaviridae* viruses contain multiple structurally conserved elements within the 5’ and 3’ untranslated regions (UTRs) that have been suggested to play roles in viral gRNA and sgRNA replication. These elements include three stem-loops (SL1, SL2, and SL3) within the 5’ UTR, as well as a bulged stem-loop (BSL), pseudoknot (PK) stem-loop, and hypervariable region (HVR) within the 3’ UTR. Recent structural studies utilizing NMR spectroscopy, SHAPE mutational profiling, and various other characterization metrologies have confirmed the conservation of these elements within the SARS-CoV-2 genome (12–14). Moreover, a recent investigation of the short- and long-range interactions within the SARS-CoV-2 genome suggests these elements additionally function in genome cyclization and interactions with the frame-shifting elements of the gRNA ORF regions, indicating potential mechanistic roles in discontinuous transcription and viral protein translation events as well (15).

While the majority of the SARS-CoV-2 3’-UTR is variable in sequence, it contains a highly conserved 41-nucleotide (nt) stem-loop II motif (s2m) within the terminal portion of the HVR (Figure 1) (16). This element has previously been shown to be conserved not only among *Coronaviridae* viruses, but also within the distantly related *Astroviridae, Caliciviridae*, and *Picornaviridae* families of positive-sense single-stranded RNA viruses (16, 17). The exact function of the s2m element has yet to be elucidated, but given its presence in several families of viruses, it has been suggested to confer a selective advantage. Proposed hypotheses for the role of the s2m element include hijacking of host protein synthesis, involvement within RNA interference pathways, and RNA recombination events (16, 18, 19). Nonetheless, the broad conservation of the s2m element in multiple families of viruses highlights the potential significance of the motif in virus detection and therapeutic drug targeting (20).

**Figure 1.**
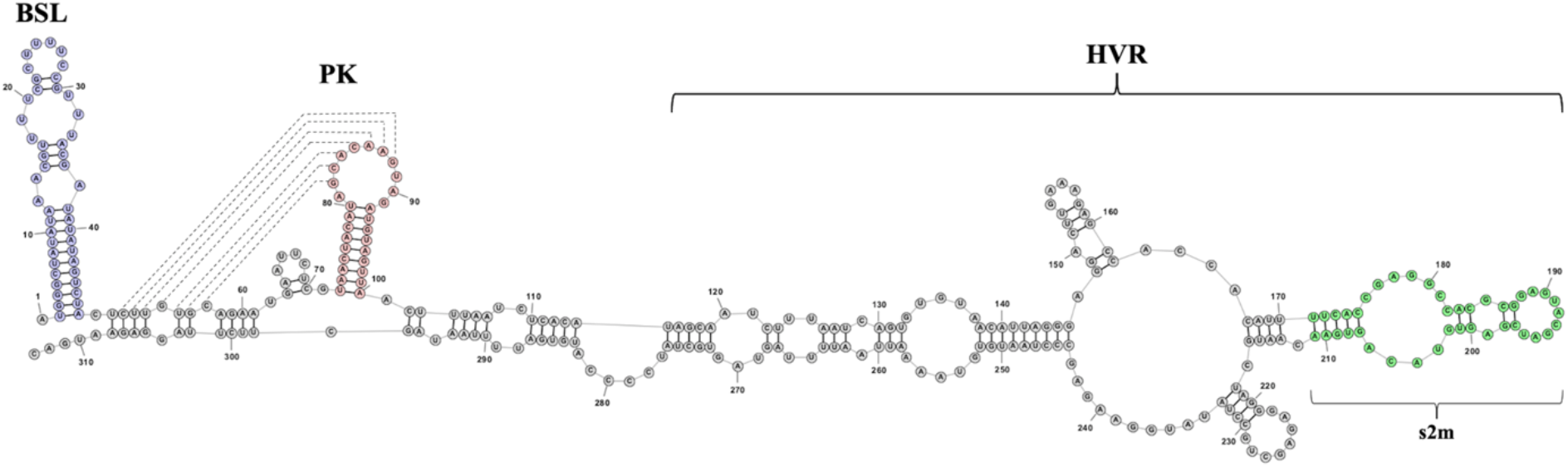
Schematic representation of the SARS-CoV-2 3’ UTR. Structure was folded using RNAstructure software and edited using StructureEditor software (56). Conserved elements are highlighted: BSL (purple), PK stem-loop (pink), and s2m element (green). Dashed lines denote pseudoknot interactions.

In this study, we identified a 4-nt palindrome (GUAC) within the terminal loop of the SARS-CoV-2 s2m element, which we show is involved in homodimeric RNA-RNA kissing complex formation. Furthermore, we show that this kissing dimer is able to undergo subsequent conversion into a thermodynamically stable duplex conformation, facilitated by the chaperone activity of the SARS-CoV-2 N protein, a process that may stabilize a dimerized form of the gRNA. This mechanism has been described previously in both hepatitis C virus (HCV) and human immunodeficiency virus 1 (HIV-1), suggesting potential roles of the s2m element in viral replication and homologous recombination events (21–24). Moreover, recent studies, as well as our own bioinformatic analysis, have identified multiple binding sites within the SARS-CoV-2 s2m element for the host miRNA-1307-3p, which has the potential to regulate the production of various interleukins and interleukin receptors that have been linked to the “cytokine storm” reported in severe COVID-19 patients (25). Given these potential host-virus RNA interactions, we further proposed that the SARS-CoV-2 s2m element is able to hijack miRNA-1307-3p for its own advantage. Our results advance the breadth of knowledge surrounding the highly conserved s2m element, as well as highlight its potential significance as a therapeutic target for both COVID-19 and other coronavirus-related diseases.

## MATERIALS AND METHODS

### Oligonucleotide Synthesis

RNA and miRNA oligonucleotides used in this study (Table 1) were chemically synthesized by Dharmacon, Inc. Lyophilized samples were re-suspended in sterile, deionized water or in 10 mM cacodylic acid, pH 6.5, prior to data acquisition.

**Table 1.**
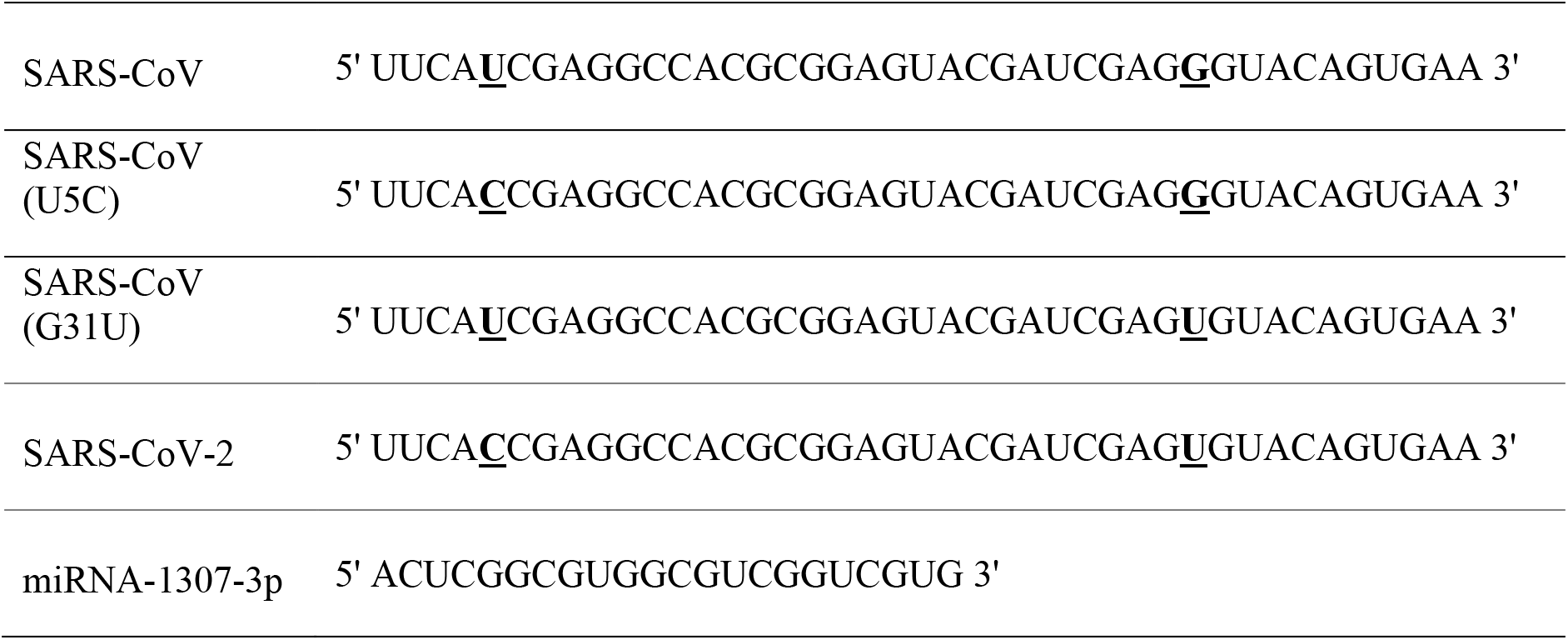
RNA sequences used in this study. Locations of nucleotide differences between SARS-CoV and SARS-CoV-2 (U5C and G31U) are highlighted in bold and underlined.

### Nondenaturing Polyacrylamide Gel Electrophoresis

Oligonucleotides were diluted in ½x Tris-Boric Acid (TB) buffer and annealed at 95 °C, followed by snap-cooling using dry ice and ethanol. To test the effect of magnesium on kissing complex formation, 500 nM or 1 μM RNA samples were incubated for either 60 minutes or 24 hours at 22 °C with increasing concentrations of MgCl_2_ in the range of 1-10 mM. Conversion of the s2m kissing complex to an extended duplex by the SARS-CoV-2 N protein (RayBiotech) was monitored by incubating 1 μM RNA samples with 1 mM MgCl_2_ and 2 μM of the protein for 45 minutes, followed by the addition of proteinase K for 15 minutes to digest the N protein. To monitor complex formation of the s2m RNA with miRNA-1307-3p, 500 nM or 1 μM RNA was incubated with 1 mM MgCl_2_ for 30 minutes, followed by addition of increasing concentrations of miRNA for an additional 30 minutes. Following incubation, samples were split and electrophoresed in Tris-Boric Acid-EDTA (TBE) or Tris-Boric Acid-Magnesium (TBM) gels for 2-hours and 4-hours, respectively. TBM gels contained 5 mM MgCl_2_ in both the gel and the ½x TB running buffer. Gels were subsequently stained with the cyanine dye SYBR® Gold and visualized by UV transillumination using an AlphaImager equipped with a 537 nm emission filter. Experiments were performed at least in duplicate.

### NMR Spectroscopy

One-dimensional ^1^H NMR spectroscopy was performed at 19 °C on a 500 MHz Bruker AVANCE NMR spectrometer equipped with TopSpin3.2 acquisition software. RNA samples at a concentration of 250 μM were prepared in 10 mM cacodylic acid, pH 6.5, in a 90:10 H_2_O:D_2_O ratio. Samples were annealed at 95 °C and snap-cooled using dry ice and ethanol prior to data acquisition. Water suppression was carried out using the Watergate pulse sequence (26).

A ^1^H-^15^N SOFAST-HMQC at ^15^N natural isotopic abundance was collected on a 900 MHz NMR AVANCE III spectrometer (Bruker BioSpin) equipped with triple resonance cryogenically-cooled probe with a z-axis gradient system. The experiment was recorded with the ^1^H and ^15^N carriers placed at 4.694 ppm and 150 ppm, respectively, with a sweep width of 22.04 ppm and 30 ppm, respectively. The recycle delay was set to 0.4 s, 25,600 scans per transient with acquisition times of 50 ms and 8.8 ms in the ^1^H and ^15^N dimensions, respectively. Selective excitation and refocusing of the amide resonances was achieved using a 90° PC-9 and 180° Reburp shaped pulses, respectively, applied at 11.5 ppm with a bandwidth of 6.0 ppm. The experiment, processed in NMRPipe, was apodized with a shifted sine-squared bell, zero-filled to quadruple the points in ^1^H and NUS-zero filled to twice the points in ^15^N using iterative soft thresholding (27). The spectrum was visualized with NMRFAM-Sparky 3.19 (28).

## RESULTS AND DISCUSSION

### The s2m region in the 3’ UTR of SARS-CoV-2 forms a homodimeric kissing complex

From the proposed SARS-CoV-2 s2m stem-loop structure, a 4-nt palindrome GUAC was observed within the loop, suggesting a possible site for the formation of a homodimeric kissing complex that could be implicated in genome dimerization (Figure 2) (12). To investigate this proposed interaction, nondenaturing gel electrophoresis studies were carried out both in the presence and absence of Mg^2+^ ions (Figure 3). Samples of the SARS-CoV-2 s2m element at concentrations of 500 nM and 1 μM were prepared in the presence of 1, 5, and 10 mM MgCl_2_, conditions which stabilize the kissing dimer conformation, and subsequently split to be run in parallel in TBE and TBM gels (29, 30). As the TBM gels contain 5 mM MgCl_2_, kissing dimer conformations are retained, whereas these ions are chelated by EDTA in the TBE gels, resulting in kissing dimer dissociation (24, 29, 30). In the TBM gel, we observed three distinct bands at all concentrations of Mg^2+^ ions investigated (Figure 3A, right), with the lower band (arrow 1) attributed to the monomeric species and the higher bands (arrows 2 and 3) to two distinct dimeric species. In contrast, upon chelation of Mg^2+^ ions by the EDTA in TBE buffer, the two dimeric bands collapse into a single band, indicating that the motif exists primarily in its monomeric state (Figure 3A, left panel, arrow 1). As has been noted previously, the s2m element monomer bands are observed to migrate differently on the TBE and TBM gels due to the effect of the Mg^2+^ ions on the monomeric structure.

**Figure 2.**
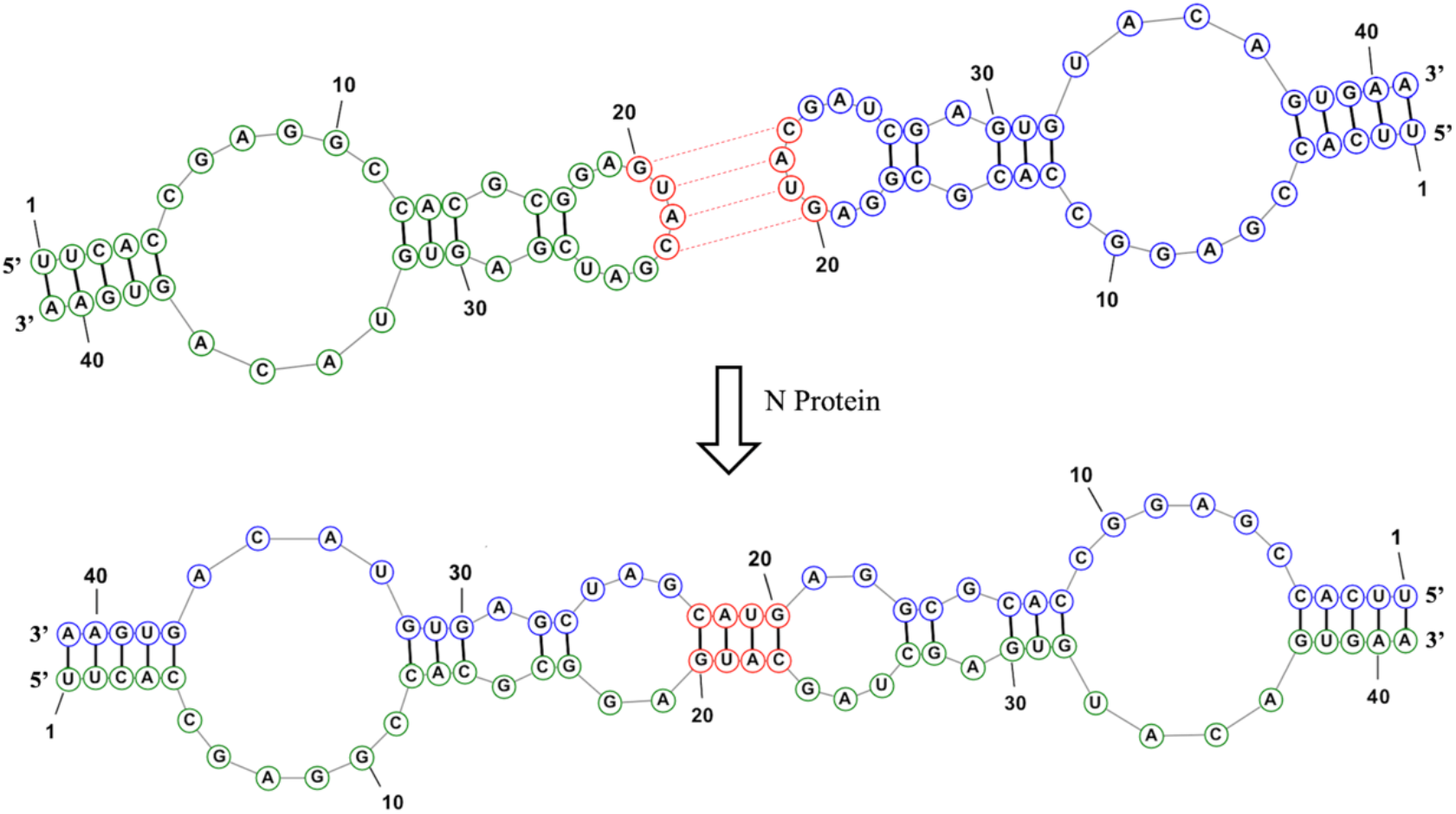
Schematic depiction of proposed s2m dimerization mechanism. Kissing complex formation is mediated by a 4-nt GUAC palindromic sequence located within the loop region of the motif. The viral N protein acts as a molecular chaperone to aid conversion of the kissing dimer conformation (top) to a thermodynamically stable extended duplex structure (bottom).

**Figure 3.**
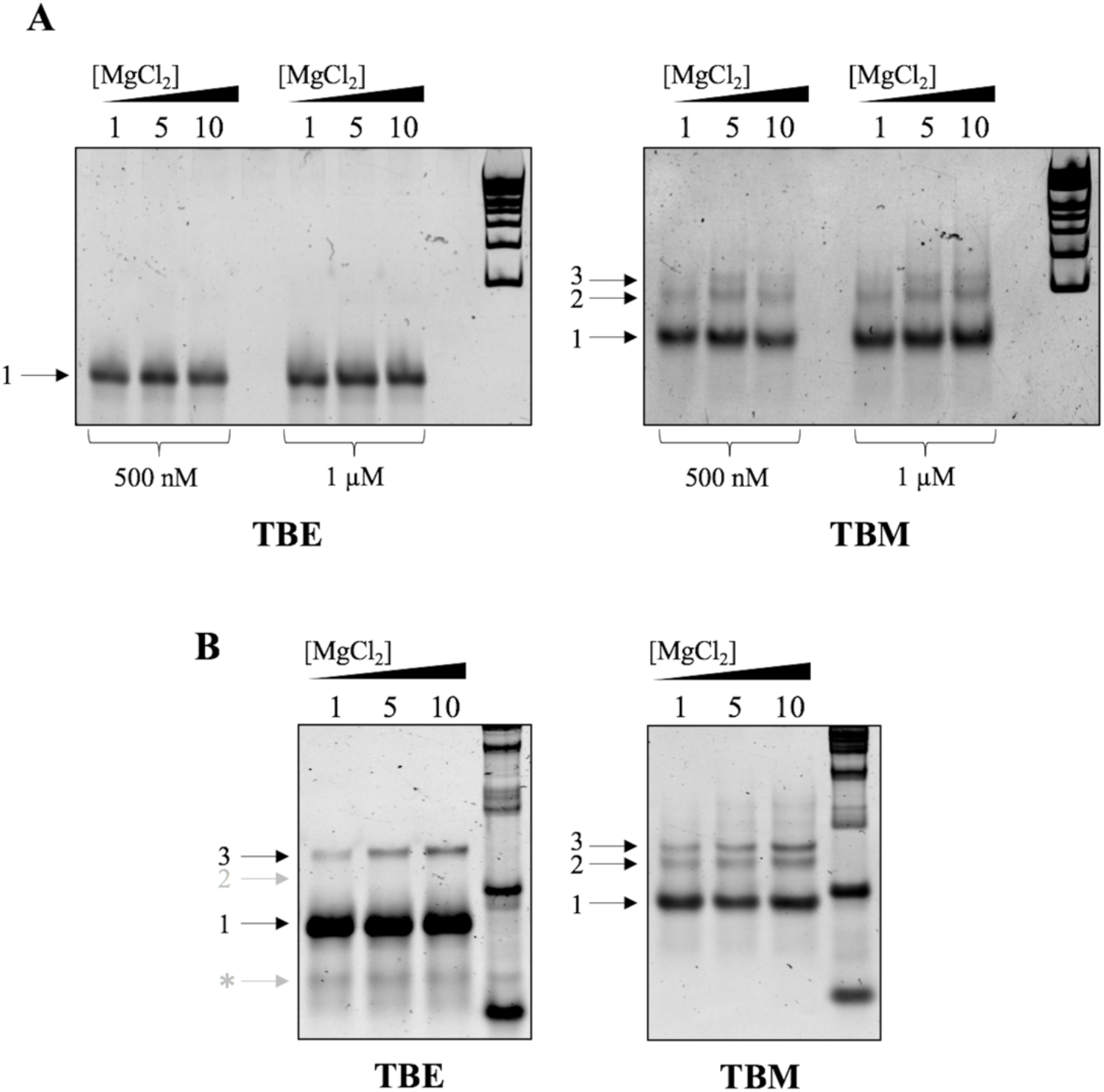
Mg^2+^-dependent nondenaturing TBE and TBM gel electrophoresis of SARS-CoV-2 s2m. Samples at RNA concentrations of 500 nM and 1 μM were incubated for 1-hour in the presence of 1, 5, and 10 mM MgCl_2_ (**A**) prior to being split and electrophoresed. In the absence of Mg^2+^ ions, the s2m element exists primarily in a monomeric conformation (left panel, arrow 1), while the presence of Mg^2+^ ions promotes the formation of both a kissing complex (arrow 2) and stable, extended duplex (arrow 3), which exists in equilibrium with the monomeric s2m (right panel). Similarly, samples at an RNA concentration of 1 μM were incubated in the presence of 1, 5, and 10 mM MgCl_2_ for 24-hours (**B**) prior to being split and electrophoresed. Extended incubation time promoted the spontaneous conversion from the kissing dimer conformation (arrow 2) to a stable, extended duplex (arrow 3) which is present even upon chelation of Mg^2+^ ions (left panel). Degradation of s2m RNA occurred during 24-hour incubation, as denoted by an asterisk (*).

We hypothesized that one of the dimer bands observed in the TBM gel corresponded to a homodimeric kissing complex, while the other arose from the spontaneous conversion of the kissing complex to a stable, extended duplex (Figure 2, bottom) which occurred during the incubation with MgCl_2_ and subsequent 4-hour run through the TBM gel. Spontaneous conversion of labile palindromic kissing dimers to their extended duplex conformation has been reported for the dimerization initiation site of other viruses, such as HCV and HIV-1 (21-24). The process, which involves strand exchange between the two monomers, is dependent on the intrinsic stability of the monomeric stem-loop structures (31, 32). The conversion of the s2m kissing dimer to duplex following the 1-hour incubation with Mg^2+^ ions was likely hindered by chelation of the ions by EDTA in the TBE gel, resulting in a band too faint to be detected. To test this hypothesis, 1 μM SARS-CoV-2 s2m samples were incubated in the presence of increasing concentrations of MgCl_2_ at room temperature for 24-hours and subsequently examined by TBE and TBM gel electrophoresis. Both dimer bands are present in the TBM gel (Figure 3B, right panel, arrows 2 and 3) with the uppermost dimer band (arrow 3) increasing in intensity at higher MgCl_2_ concentrations. Similarly, a dimer band which increases in intensity as the MgCl_2_ concentration increases is now clearly visible in the TBE gel (Figure 3B, left panel, arrow 3), indicating that the band corresponds to an extended duplex, which is stable even following chelation of Mg^2+^ ions by EDTA. Notably, both dimer bands are more intense in the TBM gel (compare Figures 3A and 3B, right panels), with the intensity of the uppermost dimer band increasing as concentration of MgCl_2_ in the sample is increased, indicating stabilization of the kissing complex intermediate followed by conversion to the duplex conformation during the 24-hour incubation time (Figure 3B). Our results support the hypothesis that the s2m element in the 3’ UTR of SARS-CoV-2 forms a homodimeric kissing complex stabilized by Mg^2+^ ions and that this structure can serve as an intermediate in the spontaneous rearrangement into a stable, extended duplex structure.

Bioinformatic analysis of the SARS-CoV-2 genome revealed the s2m element differs in sequence from that in SARS-CoV by only 2 nucleotides (U5C and G31U, Table 1). To determine whether these minor variations in the motif have an effect on its ability to dimerize, we incubated both SARS-CoV and SARS-CoV-2 s2m elements in the presence of 1, 5, and 10 mM MgCl_2_ for 1-hour, followed by electrophoresing in TBE and TBM nondenaturing gels (Figure 4). In the absence of Mg^2+^ ions, both SARS-CoV and SARS-CoV-2 s2m elements exist primarily in their monomeric state (Figure 4, left panel). Surprisingly, in the presence of Mg^2+^ ions in the TBM gel, the equilibrium is shifted more significantly towards the dimeric state for SARS-CoV s2m as compared to that of SARS-CoV-2 s2m (Figure 4, right panel, arrow 2). Additionally, while two dimer bands of lower intensity are present for the SARS-CoV-2 s2m (Figure 4, right panel, arrows 2 and 3), only a single dimeric conformation is evident for SARS-CoV (Figure 4, right panel, arrow 2). Incubation of the SARS-CoV s2m with 1, 5, and 10 mM MgCl_2_ for 24-hours at room temperature revealed no upper dimer band present in the TBE gel (Supplemental Figure S1, left panel), confirming the inability of the element to convert from the kissing dimer to the stable duplex structure. Thus, our results suggest that SARS-CoV s2m exists primarily in a kinetically trapped kissing dimer structure that does not spontaneously convert to a stable duplex. To minimize the spontaneous conversion of the SARS-CoV-2 s2m kissing dimer to duplex conformation, RNA samples were incubated in the presence of 1 mM MgCl_2_ for 1-hour in all subsequent experiments.

**Figure 4.**
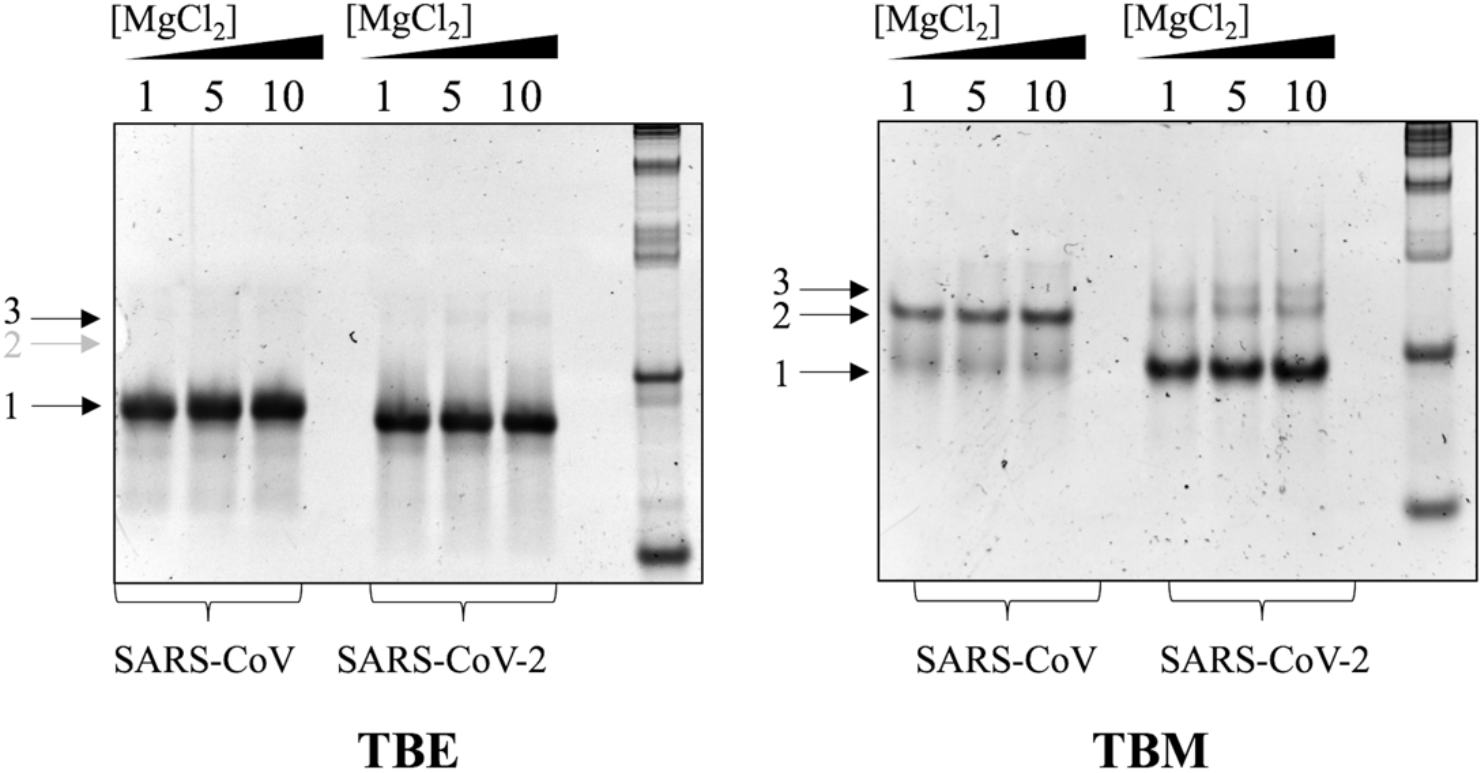
Mg^2+^-dependent comparison of SARS-CoV and SARS-CoV-2 s2m elements by nondenaturing TBE and TBM gel electrophoresis. Both s2m elements at RNA concentrations of 1 μM were incubated in the presence of 1, 5, and 10 mM MgCl_2_ for 1-hour prior to being split and electrophoresed. Both SARS-CoV and SARS-CoV-2 s2m elements exist primarily in their monomeric states upon chelation of Mg^2+^ ions in the TBE gel (left panel, arrow 1). In the presence of Mg^2+^ ions in the TBM gel, SARS-CoV exists primarily in dimeric kissing complex conformation (right panel, arrow 2), whereas SARS-CoV-2 exists in equilibrium between its monomeric (arrow 1) and two dimeric conformations (arrows 2 and 3).

The difference in dimerization properties of the s2m elements is significant, considering that the SARS-CoV-2 s2m sequence differs from that of SARS-CoV only at two positions, U5C and G31U (Table 1). The nucleotide at position 5 in the s2m element is variable; however, that at position 31 remains invariant in all viruses in which the element has been identified (16, 17, 33). According to the X-ray crystal structure of SARS-CoV s2m, U5 is located in the lower stem and base-paired to G37, whereas G31 is located in the middle stem, base-paired with C12 (Figure 5A) (18). The secondary structure of the SARS-CoV-2 s2m element was recently solved by NMR spectroscopy and is conformationally different from that of SARS-CoV s2m (12). These conformational variations were additionally confirmed by our 1.5 μs molecular dynamic simulations (manuscript submitted to Biophysical Journal, Special Issue: Biophysicists Address COVID-19 Challenges). However, it should be noted that while C5 remains base-paired to G37 and retains its position in the lower stem, U31 in SARS-CoV-2 s2m is part of a smaller middle stem, forming a base pair with A13 (Figure 5B). The terminal loop region in SARS-CoV s2m is bent into a 90° kink facilitated by a base quartet interaction, leaving G20 and U21 exposed and possibly orienting them in the right conformation for kissing dimer formation (18). In contrast, the terminal loop of SARS-CoV-2 is larger, with the entire GUAC palindrome left exposed. Selective 2’ hydroxyl acylation by primer extension (SHAPE) analysis of the s2m element in SARS-CoV-2 confirms the lability of all four nucleotides (13, 14). Thus, we postulate that the higher flexibility of the entire GUAC loop results in a greater entropic penalty upon SARS-CoV-2 s2m kissing dimer formation. These observations potentially explain why SARS-CoV s2m more easily forms a stable kissing complex structure as compared with SARS-CoV-2, which prefers its monomeric conformation even in the presence of MgCl_2_ (Figure 4, right panel).

**Figure 5.**
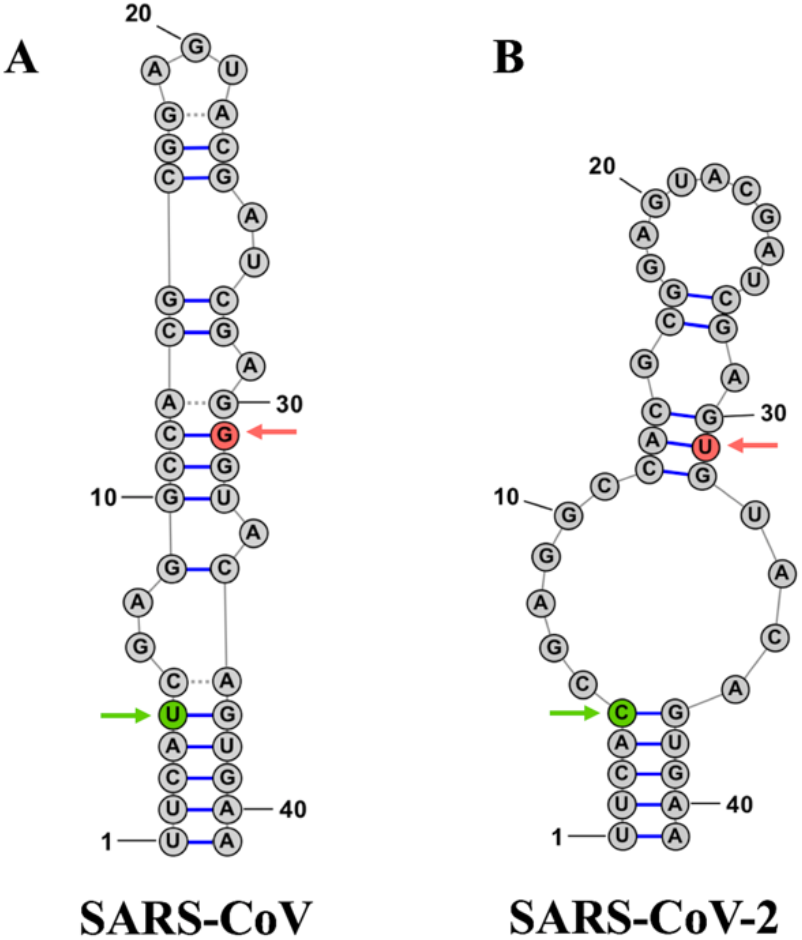
Schematic representation of SARS-CoV (**A**) and SARS-CoV-2 (**B**) s2m element secondary structures. Structures are adapted from SARS-CoV and SARS-CoV-2 s2m elements experimentally characterized by X-ray crystallography and NMR spectroscopy, respectively (12, 18). Locations of nucleotide differences are highlighted in green (U5C) and red (G31U). Structures of SARS-CoV and SARS-CoV 2 s2m elements were created using RNAstructure software and edited using StructureEditor software (56).

To determine whether the structural differences and altered dimerization properties of SARS-CoV-2 s2m are due to both nucleotide variations, we investigated two mutated SARS-CoV sequences in which a single nucleotide mutation was introduced at a time (U5C and G31U, respectively, Table 1 and Figure 5). We employed one-dimensional ^1^H NMR spectroscopy to analyze the secondary structures of these mutants to determine if they resemble the SARS-CoV s2m or SARS-CoV-2 s2m secondary structure. The SARS-CoV-2 s2m imino proton resonance region (Figure 6A, top spectrum) matched that of its previously determined secondary structure, allowing us to assign its resonances (12). It should be noted, however, that our analysis focused on a synthesized 41-nucleotide sequence corresponding to the SARS-CoV-2 s2m element, whereas the previous characterization included two additional G-C base pairs introduced in the lower stem for transcriptional purposes. In the SARS-CoV (G31U) s2m spectrum, the imino proton resonances corresponding to U31, G30, G32, and G28 have similar chemical shifts as compared with the corresponding ones in SARS-CoV-2 s2m (Figure 6A). However, those resonances corresponding to base pairs in the lower stem appear to be broader, potentially due to the fact that in wild-type SARS CoV, the G-C base pair formed by C5-G37 is replaced by the less stable U5-G37 (compare Figure 5). As expected, the U31 imino proton resonance is absent from both wild-type SARS-CoV s2m and SARS-CoV (U5C) s2m mutant, and while we cannot make unambiguous assignments for these imino proton resonances, we noted that the overall spectra corresponding to wild-type SARS-CoV s2m and its U5C mutant appear to be similar (Figure 6A, bottom two spectra). We noted that the imino proton resonances for both SARS-CoV s2m and its U5C mutant are broader than those for SARS-CoV-2 s2m and SARS-CoV (G31U), which could be due to the formation of kissing dimer structures even in the absence of Mg^2+^ ions at the high RNA concentrations used in NMR spectroscopy experiments.

**Figure 6.**
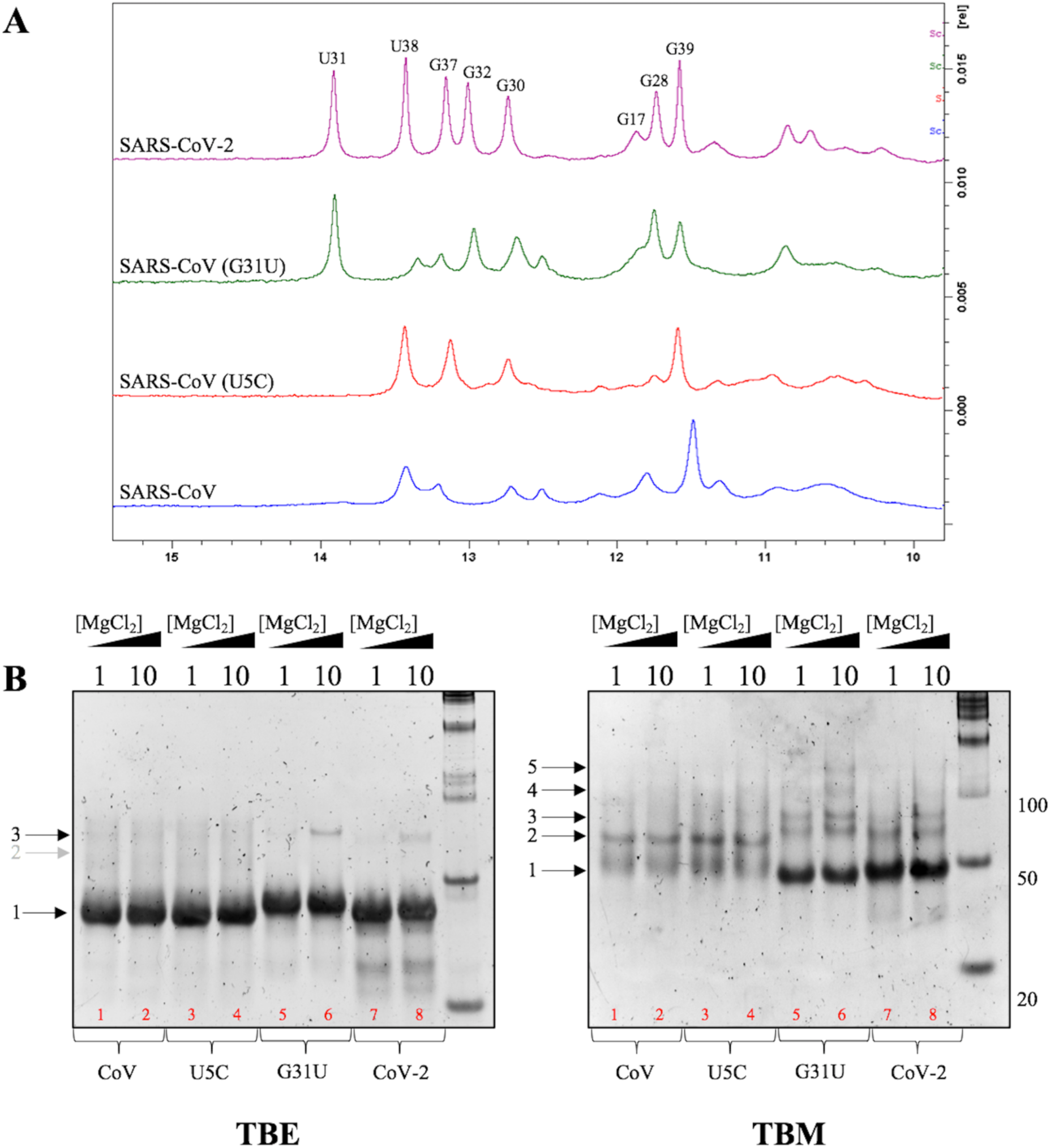
Characterization of the s2m element mutant constructs and their dimerization properties. One-dimensional ^1^H NMR spectroscopy experiments were performed for all four s2m constructs in the presence of 0 mM MgCl_2_ (**A**), with resonances being assigned within the SARS-CoV-2 s2m spectrum based on the previously characterized secondary structure (12). All four s2m sequences were additionally incubated in the presence of 1 and 10 mM MgCl_2_, followed by electrophoresing in TBE and TBM nondenaturing gels (**B**). In the presence of Mg^2+^ ions (right panel), wild-type SARS-CoV and SARS-CoV (U5C) constructs exist primarily in a kissing dimer conformation (arrow 2), whereas SARS-CoV (G31U) and SARS-CoV-2 s2m constructs exist in equilibrium between monomeric (arrow 1) and two dimeric conformations (arrows 2 and 3). Higher-order complexes within SARS-CoV (G31U) s2m are denoted by arrows 4 and 5.

Based on these ^1^H NMR spectroscopy results, we predicted that the dimerization of the SARS-CoV (G31U) s2m mutant will be similar to that of wild-type SARS-CoV-2 s2m, whereas that of the SARS-CoV (U5C) s2m mutant will resemble that of wild-type SARS-CoV s2m. Thus, we performed nondenaturing PAGE for these mutants in the presence of 1 and 10 mM MgCl_2_. In the TBM gel (Figure 6B, right panel), the SARS-CoV (U5C) s2m mutant, like the wild-type SARS-CoV s2m, shows a single dimer band (arrow 2). In contrast, the SARS-CoV (G31U) s2m mutant, like SARS-CoV-2 s2m, shows two distinct, albeit more intense, dimer bands which correspond to kissing dimer (arrow 2) and extended duplex (arrow 3). Additional higher molecular weight bands above the 100-bp marker are also evident for the SARS-CoV (G31U) s2m at 10 mM MgCl_2_, indicating the formation of higher-order complexes for this mutant (Figure 6B, right panel, arrows 4 and 5).

Upon chelation of Mg^2+^ ions in the TBE gel, all four s2m elements exist primarily in their monomeric conformation (Figure 6B, left panel). However, a distinct dimer band is also apparent for the SARS-CoV (G31U) s2m mutant at both 1 and 10 mM MgCl_2_ (Figure 6B, left panel, arrow 2), indicating that this mutant kissing dimer converts more readily to the stable, extended duplex conformation, as compared to SARS-CoV-2 s2m. This is likely due to a less stable lower stem, which is capped by the U5-G37 base pair in SARS-CoV (G31U) s2m, as compared with the C5-G37 base pair in the wild-type SARS-CoV-2 s2m. These results, which also corroborate the ^1^H NMR data, indicate similar dimerization properties between wild-type SARS-CoV s2m and SARS-CoV (U5C) s2m mutant (Figure 6A, bottom two spectra), as both exist primarily in a kissing dimer conformation in the presence of Mg^2+^ ions (Figure 6B, right panel, lanes 1-4).

Taken together, these results indicate that the different nucleotide at position 31, a previously 100% conserved G in the s2m element of SARS-CoV which is mutated to U in the SARS-CoV-2 s2m, is the single driving force in the drastic difference in dimerization properties of this motif. To gain additional higher resolution information about the U31 nucleotide in SARS-CoV-2 s2m, we performed ^1^H-^15^N SOFAST-HMQC NMR spectroscopy experiments on a 900 MHz spectrometer. Although the U31 imino proton gives rise to a single, sharp resonance in the one-dimensional ^1^H spectrum of SARS-CoV-2 s2m acquired at 500 MHz (Figure 6A, top spectrum), three distinct cross peaks are apparent in the two-dimensional ^1^H,^15^N SOFAST-HMQC spectrum acquired at 900 MHz (Supplemental Figure S2A), indicating that this position is conformationally dynamic. This could lead to the destabilization of the middle stem region of the s2m element in SARS-CoV-2, allowing it to more readily convert to the extended duplex conformation (Figure 6B) as compared to SARS-CoV s2m.

### The viral N protein converts the SARS-CoV-2 s2m kissing dimer to a stable, extended duplex

During the viral life cycle of coronaviruses, the N protein is primarily responsible for packaging the viral RNA(+) genome into compact ribonucleoprotein (RNP) complexes during virion assembly (34, 35). Previous work on HIV-1 and HCV has elucidated additional roles for the nucleocapsid protein, demonstrating that it acts as a molecular chaperone, converting kissing complex formations into stable, extended duplexes (23, 24, 36–38). Given our finding that the s2m element of SARS-CoV and SARS-CoV-2 form homodimeric kissing complexes, we proposed a similar chaperoning ability by the N protein. To test this hypothesis, nondenaturing TBE and TBM gel electrophoresis was employed to monitor the formation of a thermodynamically stable duplex, as previously described (23, 24).

Both SARS-CoV and SARS-CoV-2 s2m elements were incubated in the presence of 1 mM MgCl_2_ to pre-form kissing complex structures, followed by addition of the SARS-CoV-2 N protein in a 1:2 ratio and incubation for an additional 30-minutes. To monitor chaperone activity, proteinase K was subsequently added to the samples to degrade the N protein prior to electrophoresing. Upon chelation of Mg^2+^ ions in the TBE gel, a distinct dimer band was present in SARS-CoV-2 s2m, indicating the formation of a stable duplex (Figure 7, left panel, arrow 3). Surprisingly, no distinct dimer band was present in the SARS-CoV s2m sample incubated with the N protein. In order to ensure the observed effect was not due to non-specific interactions, both RNAs were incubated with 2 μM bovine serum albumin (BSA) under similar conditions, and in both SARS-CoV and SARS-CoV-2 s2m samples, the BSA protein had no effect on duplex formation (Figure 7, left panel). The increased intensity of the top dimer band of SARS-CoV-2 s2m after the action of the N protein (Figure 7, right panel, arrow 3) and concomitant decrease in the monomer band (Figure 7, right panel, arrow 1) in the TBM gel confirm our previously established hypothesis that an equilibrium exists between the monomer and two dimer conformations, with the N protein shifting the equilibrium towards the stable duplex structure. Additional time-dependent studies monitoring the chaperone activity of the N protein revealed no change in duplex band intensity after incubation for 1.5-hours for SARS-CoV-2 and no apparent duplex band for SARS-CoV s2m (Supplemental Figure S3, arrow 3). To determine if the N protein has an effect on the SARS-CoV s2m conversion from the kissing dimer to duplex conformation at physiological temperatures, we performed studies at 37 °C. Similar to the results seen in Figure 7, no distinct dimer band was present for SARS-CoV s2m following incubation with the N protein (Supplemental Figure S4, left panel, arrow 3), but the higher temperature accelerated the spontaneous conversion from kissing complex to duplex conformations for SARS-CoV-2 s2m, as bands corresponding to extended duplex are clearly apparent even in the absence of the N protein (Supplemental Figure S4, left panel, arrow 3).

**Figure 7.**
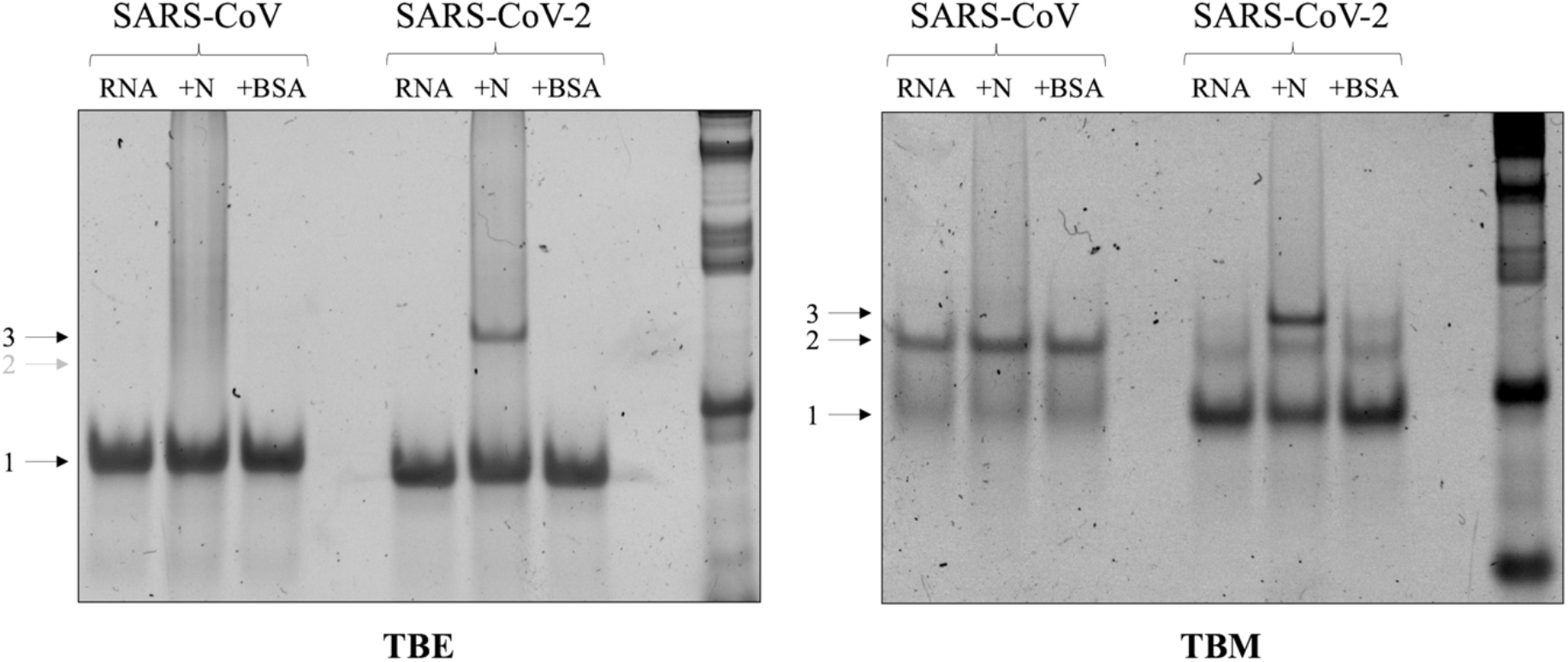
Conversion of the SARS-CoV-2 s2m kissing complex to a stable, extended duplex by the viral N protein. Both SARS-CoV and SARS-CoV-2 s2m elements were incubated in the presence of 1 mM MgCl_2_ for 1-hour, followed by addition of the N protein in a 1:2 ratio and additional incubation for 30-minutes at 22 °C. Proteinase K was added to digest the N protein prior to samples being split and electrophoresed by TBE and TBM nondenaturing PAGE. The kissing complex formed by SARS-CoV-2 s2m (arrow 2) is converted to a thermodynamically stable extended duplex (arrow 3) by chaperone activity of the N protein, whereas SARS-CoV s2m is unable to undergo similar conversion. Control samples in which both s2m elements were incubated in the presence of BSA in a 1:2 ratio revealed no changes in conversion to stable duplex conformation.

The finding that the SARS-CoV s2m kissing dimer is not converted to a stable duplex conformation is surprising considering that the two sequences differ only by two nucleotides (Table 1 and Figure 5). In these experiments, we used the SARS-CoV-2 N protein, whose RNA-binding domain has a high sequence and structural homology with that of the SARS-CoV N protein (Supplemental Figure S5). Nonetheless, we performed nondenaturing PAGE in the absence of proteinase K to determine if the SARS-CoV-2 N protein can bind to both the SARS-CoV and SARS-CoV-2 s2m elements, and as seen in Figure 8 (arrows 2-4), the protein binds both RNA sequences. We further demonstrated in control experiments that the N protein binds specifically to these particular RNA stem-loop structures, as it did not bind to the SARS-CoV-2 5’ UTR SL2 and SL3 stem-loops and bound weakly to the SARS-CoV-2 3’ UTR pseudoknot (PK) stem-loop (Supplemental Figure S6). Thus, we propose that the lack of conversion from the kissing dimer to a duplex conformation in SARS-CoV s2m is due to the fact that its structure lacks the large internal bulge (compare Figures 5A and 5B), which makes it too stable for N protein unfolding. Previous studies of the HIV-1 nucleocapsid protein chaperone activity suggested that the presence of bulges in the lower stem of the dimer initiation site monomeric stem-loop is essential for the conversion of the kissing dimer to duplex conformation (39, 40).

**Figure 8.**
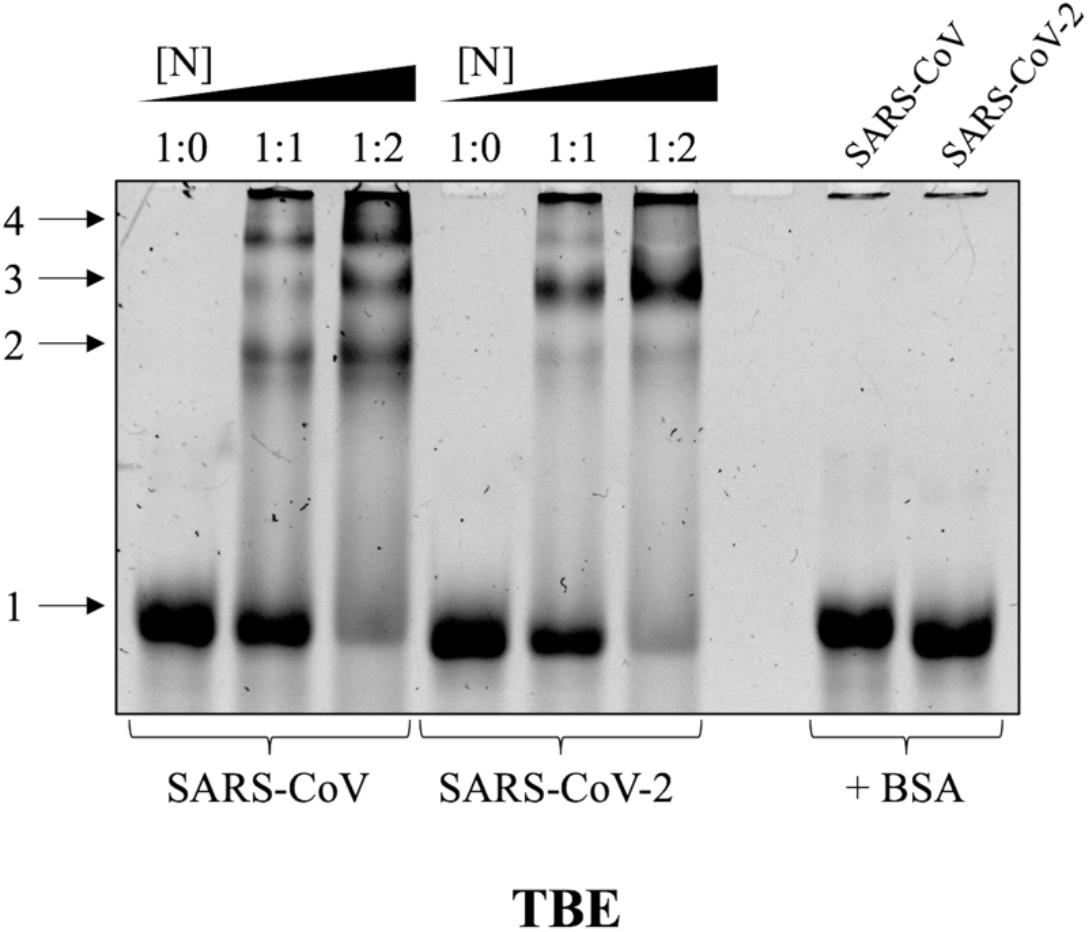
Binding of both s2m elements to the viral SARS-CoV-2 N protein. SARS-CoV and SARS-CoV-2 s2m elements were incubated in the presence of 1 mM MgCl_2_ and the SARS-CoV-2 N protein in a 1:2 ratio for 1-hour, followed by electrophoresing in a TBE nondenaturing gel. The formation of s2m:N protein complex bands (arrows 2-4) was apparent with a concomitant decrease in s2m monomeric bands (arrow 1) for both s2m elements. Control samples of each s2m element incubated in the presence of BSA at a 1:2 ratio revealed no binding, indicating the s2m:N protein complexing was protein-specific.

### The s2m element of SARS-CoV-2 interacts with cellular miRNA-1307-3p

In recent years, it has become increasing evident that coronaviruses utilize various host cell components for their own replicative benefit (6, 41, 42). Previous studies have shown that RNA viruses may also harbour the ability to hijack the functions of host cellular miRNAs to their advantage. The HCV virus, for example, sequesters miRNA-122 in order to stabilize its RNA and facilitate replication (43, 44). Similarly, hijacking of miRNA-142-3p by the North American eastern equine encephalitis virus (EEEV) results in repression of innate immune responses and subsequently, increased severity of disease (45). In most circumstances, the location of naturally-occurring miRNA-binding sites occurs within the 3’ UTR of viral RNA genomes (46).

The bioinformatics analysis of the SARS-CoV-2 genome by us and others has revealed various potential miRNA-binding sites within both the 5’ and 3’ UTRs, including two potential binding sites for miRNA-1307-3p within the s2m element (47–49). Under normal cellular conditions, miRNA-1307-3p is predicted to regulate the translation of various interleukins (IL18, CCLS), interleukin receptors (IL6R, IL10RA, IL10RB, IL2RB, IL17RA, IL12RB2), and interferon alpha receptor (IFNAR). Upregulation of these factors have been linked to the onset of the “cytokine storm” found in severe COVID-19 patients and thought to be involved in the development of acute respiratory distress syndrome (ARDS) (25, 50, 51). ARDS is caused by immuno-inflammatory injury to the alveolar-capillary membrane, which forms the blood-air barrier in the lungs. Alveolar type II (ATII) epithelial cells are the primary targets of SARS-CoV-2 as they express the angiotensin-converting enzyme 2 (ACE2) receptors on their surface, which are bound by the S protein (52). Interestingly, ATII cells have been shown to have immunomodulatory response properties. They are known to secrete IL6, IL10, and IL18, as well as various cytokine receptors (53–55). Given the potential relevance of miRNA-1307-3p regulating these various elements of the “cytokine storm” within ATII cells, as well as the potential for multiple binding sites within the s2m element, we sought to investigate their interactions, and moreover, to determine if the dimerization of the s2m element affects these interactions.

Binding studies were completed for SARS-CoV and SARS-CoV-2 using nondenaturing TBE and TBM gel electrophoresis (Figure 9). Samples were incubated in the presence of 1 mM MgCl_2_ and increasing ratios of s2m RNA:miRNA-1307-3p for 1-hour at room temperature, after which they were separated and electrophoresed in TBE (Figure 9A) and TBM (Figure 9B) nondenaturing gels. Two upper bands (Figure 9A, left panel, arrows 4 and 5) that increase in intensity with the concomitant decrease intensity of the monomer s2m band (Figure 9A, left panel, arrow 2) are present at all miRNA-1307-3p concentrations investigated in the SARS-CoV-2 TBE gel. We assign these bands to the complex of the SARS-CoV-2 s2m element with one (arrow 4) and two (arrow 5) miRNA-1307-3p molecules, respectively. The free miRNA-1307-3p migrates as a mixture of monomer (22-nt) and dimer (44-nt) (Figure 9A, both panels, arrows 1 and 3). Nonetheless, the bands corresponding to the monomeric SARS-CoV-2 s2m (41-nt) (arrow 2) and the dimeric miRNA-1307-3p (44-nt) (arow 3) are clearly distinguishable, as with increasing concentrations of miRNA-1307-3p, the monomeric SARS-CoV-2 s2m band decreases in intensity upon their complex formation (Figure 9A, left panel). Similarly, SARS-CoV s2m also forms a complex with two miRNA-1307-3p molecules, as a clear upper band is present in its TBE gel performed in the presence of increasing miRNA concentrations (Figure 9A, right panel, arrow 5). The band corresponding to the complex formed by SARS-CoV s2m and a single miRNA-1307-3p molecule (Figure 9A, right panel, arrow 4) is not clearly visible in this gel, as the studies were performed at a constant RNA concentration of 500 nM, and we assumed it is under the limit of detection. This band becomes visible when a similar TBE gel was run at 1 μM RNA with a 1:1 and 1:2 ratio of miRNA-1307-3p for both SARS-CoV and SARS-CoV-2 (Supplemental Figure S7, arrow 4). Notably, the bands corresponding to the complexes formed by SARS-CoV s2m with miRNA-1307-3p are less intense compared to those of SARS-CoV-2 (Figure 9A, arrows 4 and 5). This results in the inability to distinguish between the dimeric miRNA-1307-3p and the monomeric SARS-CoV s2m (Figure 9A, right, arrows 2 and 3).

**Figure 9.**
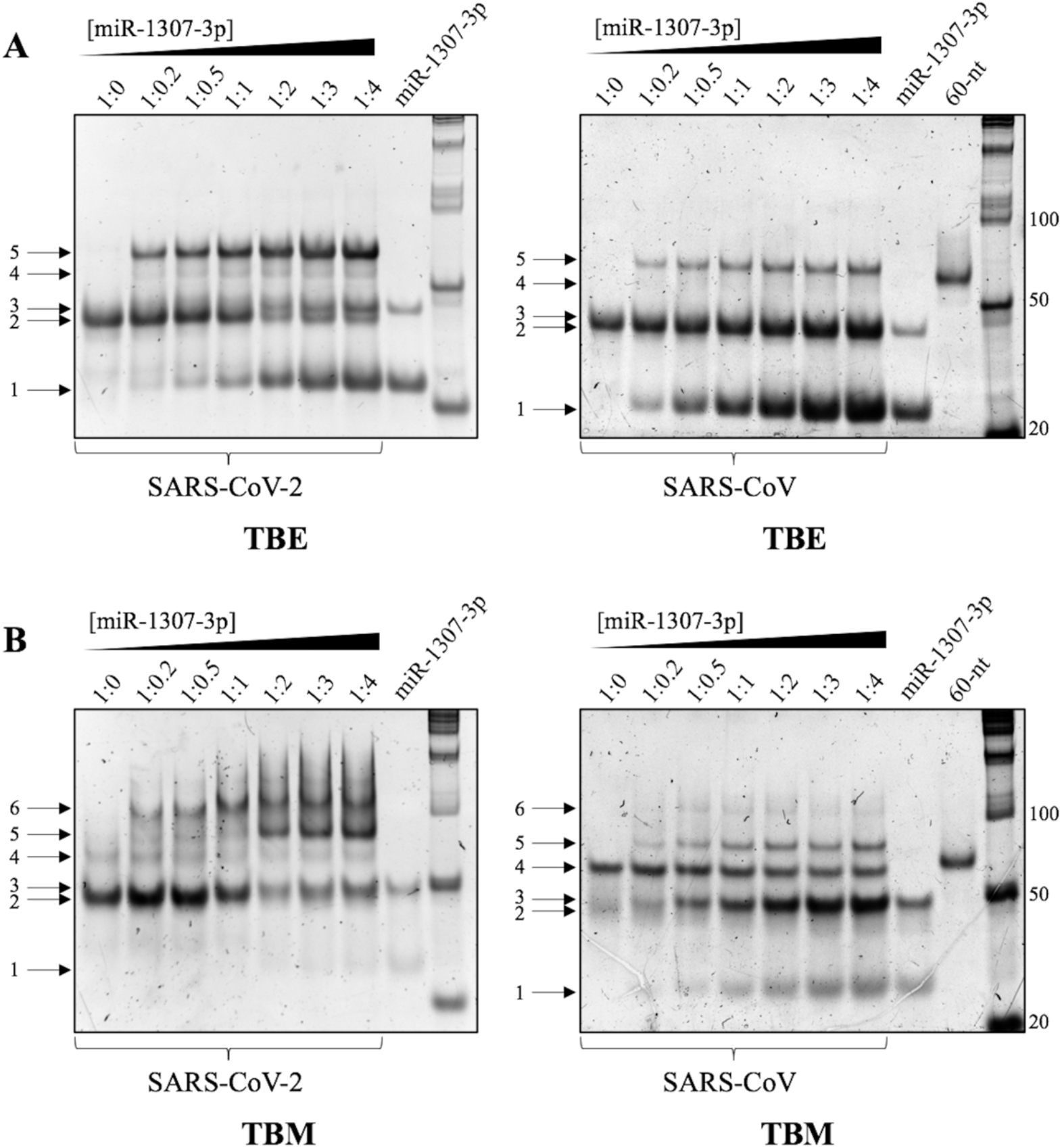
Interaction of the SARS-CoV and SARS-CoV-2 s2m elements with miRNA-1307-3p. Both s2m elements at concentrations of 500 nM were incubated with 1 mM MgCl_2_ and miRNA-1307-3p at increasing ratios (up to 1:4) for 1-hour. Samples were split and analyzed by TBE and TBM nondenaturing PAGE. Upon chelation of Mg^2+^ ions in the TBE gel (**A**), both SARS-CoV-2 (A, left panel) and SARS-CoV (A, right panel) s2m elements bind to one (arrow 4) and two (arrow 5) molecules of miRNA-1307-3p. Both complex bands of SARS-CoV-2 increase in intensity with a concomitant decrease of the band corresponding to monomeric s2m (arrow 2). In the presence of Mg^2+^ ions in the TBM gel (**B**), both SARS-CoV and SARS-CoV-2 exist in equilibrium between their monomeric and dimeric conformations, with each able to form kissing dimer conformations (arrow 4) that can bind to one (arrow 5) or two (arrow 6) molecules of miRNA-1307-3p. In all gels, miRNA-1307-3p migrates in both monomeric (arrow 1) and dimeric (arrow 3) conformations.

The interpretation of the TBM gels is more complex since they contain Mg^2+^ ions which stabilize the dimeric conformations of both SARS-CoV and SARS-CoV-2 s2m elements (Figure 9B). The kissing dimer bands in both s2m elements are visible even upon incubation with miRNA-1307-3p up to 1:4 ratios (Figure 9B, both, arrow 4). The band corresponding to the monomeric conformation of SARS-CoV-2 s2m (Figure 9B, left panel, arrow 2) decreases in intensity as the concentration of the miRNA is increased, whereas for SARS-CoV s2m, like in the TBE gel, this is not as clear to distinguish as the s2m monomer band (Figure 9B, right, arrow 2) is overlapping with that of the miRNA-1307-3p dimer band (arrow 3). These results indicate either increased propensity of miRNA-1307-3p binding to the s2m monomeric conformations or constant equilibria shift between monomeric and dimeric conformations upon miRNA binding. We observed two distinct bands for the SARS-CoV-2 s2m RNA:miRNA-1307-3p complexes, with one migrating under and the second migrating above the 100-nt marker band (Figure 9B, left panel, arrows 5 and 6). Similar to the main complex band we observed in the TBE gel (Figure 9A, left panel, arrow 5), we assign the band indicated by arrow 5 (Figure 9B, left panel) to the complex formed by one s2m element and two miRNA-1307-3p molecules. In contrast to the TBE gel, where Mg^2+^ ions present during incubation with miRNA-1307-3p are chelated by EDTA, the Mg^2+^ ions remain present in the TBM gel, allowing the s2m element to dimerize even after binding to one miRNA-1307-3p molecule, as in this complex the palindromic sequence in the terminal loop becomes fully exposed. Thus, we assign the higher molecular weight band we observe in the TBM gel (Figure 9B, left panel, arrow 6) to the dimer formed by two s2m elements, each of which is additionally bound to a single miRNA-1307-3p molecule. The s2m conformation in which two miRNA-1307-3p molecules are bound cannot dimerize since its palindromic sequence is engaged in interactions with the second molecule of miRNA-1307-3p. Like in the TBE gels, we noted that the s2m RNA:miRNA-1307-3p complex bands have higher intensity in the case of SARS-CoV-2 (Figure 9B, left panel, arrows 5 and 6) compared to those in SARS-CoV (Figure 9B, right panel, arrows 5 and 6). Interestingly, the band corresponding to the SARS-CoV s2m kissing complex with each element bound by a single miRNA-1307-3p molecule (Figure 9B, right panel, arrow 6) is barely visible.

These differences in the binding of the SARS-CoV and SARS-CoV-2 s2m elements to miRNA-1307-3p cannot easily be explained based on the two-nucleotide difference between the sequences. The first binding site for miRNA-1307-3p in SARS-CoV has a single nucleotide difference from that of SARS-CoV-2, which results in a G:U base pair being replaced with a G:C base pair in its complex with the miRNA (U5C, Figure 5), and a predicted difference in the binding free energy of only 2.6 kcal/mol as determined by the RNAstructure software (56). The second binding site for miRNA-1307-3p differs in the nucleotide at position 31, G in SARS-CoV and U in SARS-CoV-2. However, given its location in an internal bulge, the predicted binding free energy is the same for both s2m elements and is not expected to make a difference in miRNA-1307-3p binding at this site. Thus, we hypothesized that the difference in miRNA-1307-3p binding between SARS-CoV and SARS-CoV-2 s2m elements is due to their different dimerization properties. We proposed that, when in its monomeric conformation, SARS-CoV and SARS-CoV-2 s2m elements bind similarly to miRNA-1307-3p, but that these interactions are inhibited by the s2m kissing dimer formation.

To test this hypothesis, both SARS-CoV and SARS-CoV-2 s2m samples were incubated with increasing concentrations of miRNA-1307-3p in the absence of MgCl_2_, where both elements exist in their monomeric state, followed by electrophoresing on a TBE gel (Figure 10, lanes 1-6). For comparison, two samples were incubated in the presence of 1 mM MgCl_2_, conditions in which SARS-CoV s2m exists mostly as a kissing dimer, whereas SARS-CoV-2 s2m is mostly monomeric (Figure 4, right panel) prior to the electrophoresing in the TBE gel where the Mg^2+^ ions are chelated by EDTA. As predicted, the bands corresponding to the s2m element complex formed with two miRNA-1307-3p molecules (Figure 10, arrow 5) have similar intensities, indicating that the affinity for miRNA-1307-3p does not differ between the two monomeric s2m elements (Figure 10, arrow 5, compare lanes 2 and 3 with lanes 5 and 6). However, similar to Figure 9A, in the presence of Mg^2+^ ions, the band corresponding to the SARS-CoV s2m complex with two miRNA-1307-3p molecules is clearly less intense than that of SARS-CoV-2 s2m (Figure 10, compare lanes 8 and 9). We attribute these differences to the equilibria that exists between monomeric and dimeric conformations of each s2m element in the presence of Mg^2+^ ions (Figure 4, right panel). SARS-CoV-2 s2m exists primarily in its monomeric state, allowing for increased binding to miRNA-1307-3p. In contrast, SARS-CoV s2m exists primarily in a kissing dimer complex in the presence of Mg^2+^ ions, presumably inhibiting its interactions with miRNA-1307-3p. Nonetheless, the presence of Mg^2+^ ions during the 1-hour incubation promotes the formation of the complexes between the monomeric s2m and miRNA-1307-3p, as indicated by the increased complex band intensities for both SARS-CoV and SARS-CoV-2 s2m elements as compared to those incubated in the absence of any MgCl_2_ (Figure 10, arrow 5, compare lanes 1-6 with lanes 8 and 9). Upon chelation of these ions by EDTA in the TBE gel, both s2m elements dissociate to become fully monomeric, but the monomeric miRNA-1307-3p (22-nt) is no longer available for binding as it will migrate faster through the gel. Thus, we attribute the difference in intensity between the SARS-CoV-2 and SARS-CoV s2m:miRNA-1307-3p complex bands (Figure 10, lanes 8 and 9, arrow 5) to the fact that during the their incubation in the presence of Mg^2+^ ions, SARS-CoV s2m was mostly dimeric, which inhibited its binding to miRNA-1307-3p. We conclude from these results that the s2m dimerization affects its interactions with miRNA-1307-3p.

**Figure 10.**
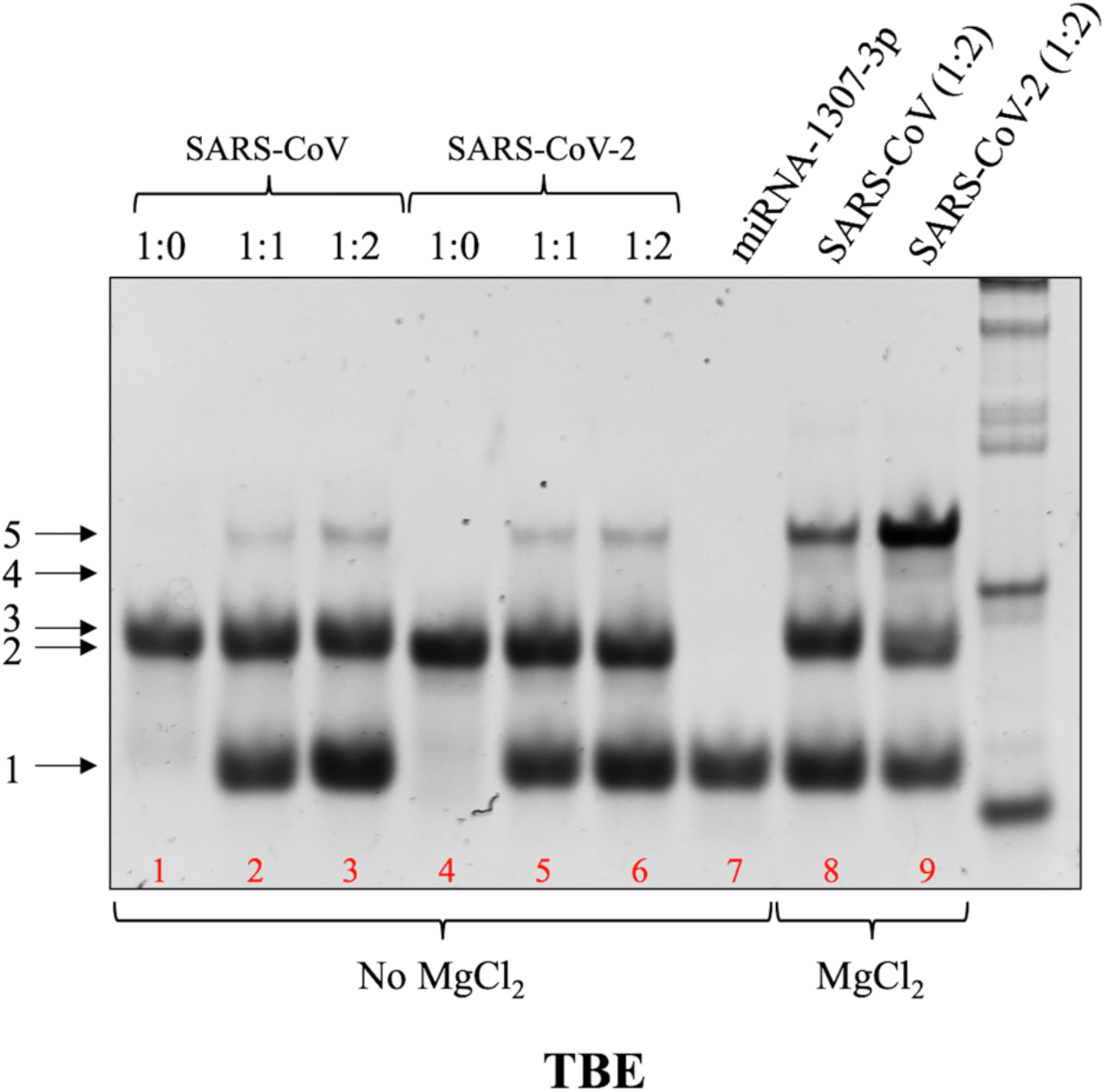
Dimerization of the s2m element affects miRNA-binding ability. SARS-CoV and SARS-CoV-2 s2m elements were incubated in the presence of miRNA-1307-3p at 1:0, 1:1, and 1:2 ratios in the absence of MgCl_2_ and electrophoresed by nondenaturing TBE PAGE. Binding affinity to the miRNA does not appear to differ between the two s2m elements, as indicated by similar complex band intensities (lanes 1-6, arrows 4 and 5). Control samples in which each s2m element was incubated with 1:2 miRNA-1307-3p in the presence of 1 mM MgCl_2_ resulted in increased complex band intensity for SARS-CoV-2 s2m compared to SARS-CoV s2m, suggesting dimerization affects miRNA-binding ability. In the absence of MgCl_2_, miRNA-1307-3p exists solely in its monomeric conformation (lane 7, arrow 1), whereas incubation with 1 mM MgCl_2_ results in the formation of a dimeric conformation as well (lanes 8 and 9, arrow 3).

While the specific role of the s2m element in coronaviruses remains unresolved, it has been proposed to be involved in viral replication, hijacking of host protein synthesis, and RNA interference pathways (16, 18, 57). Here, we demonstrate for the first time the ability of the isolated SARS-CoV-2 s2m element to form an intermediate kissing dimer which is converted to a stable duplex conformation. Moreover, our results suggest that while the SARS-CoV-2 s2m element may spontaneously convert to the thermodynamically stable duplex over time, the viral N protein exhibits chaperone activity to accelerate this conversion. In contrast, while the SARS-CoV s2m element readily forms kissing dimers, these complexes are not converted to the stable duplex conformation, even in the presence of the N protein. The conversion mechanism discovered here for SARS-CoV-2 s2m is similar to that seen in HIV-1, in which viral N protein acts as a molecular chaperone to convert transient kissing dimer complexes into a stable, extended duplex structures (22, 38, 58). This process in the *Retroviridae* family of viruses, to which HIV-1 belongs, is believed to facilitate dimerization of two genomic RNA copies for packaging in the virion, a function which does not occur in *Coronaviridae* viruses. However, an identical mechanism of monomeric stem-loop destabilization and conversion to thermodynamically stable duplex structures by the N protein has been demonstrated for HCV, another single-stranded, positive-sense RNA virus (24, 59, 60). Furthermore, the mechanism in HCV was proposed to modulate the interactions of the dimer initiation site with additional open reading frame stem-loops, which are essential for RNA replication, suggesting similar functions within SARS-CoV-2 (61, 62). It is possible that in SARS-CoV-2, this dimerization mechanism may also function in genome frameshifting and subsequent sgRNA and gRNA synthesis. A three-stemmed pseudoknot structure within the open reading frame (ORF) of the SARS-CoV genome has been shown previously to act as a ribosomal frameshift stimulating signal (63, 64). One of the stem-loop structures within this signal contains a 6-nt palindrome within its terminal loop region which has been demonstrated to form homodimeric kissing interactions, and disruption of these interactions resulted in reduced frameshifting (65). Thus, it is possible that the s2m dimerization may facilitate the alignment of multiple gRNA molecules, in coordination with the kissing loop interactions in the ORF, to allow for proper frameshifting and RNA synthesis. Lastly, we speculate on the role of genome dimerization with respect to recombination events. The origin of the SARS-CoV-2 virus is proposed to be the result of homologous recombination between bat and pangolin coronaviruses, likely triggering the cross-species transmission to humans (3, 4). High recombination rates among coronaviruses have been well documented, though the exact mechanism by which this occurs is unknown (5). Similarly, this high rate of recombination has been studied extensively in other viral families, such as *Astroviridae, Caliciviridae*, and *Picornaviridae* (66–68). Interestingly, all of these viral families also contain the conserved s2m element within their 3’ UTR (16, 19, 69). Given the high degree of similarities between recombination events and conservation of the s2m element within these additional viral families, we speculate whether this element may be involved in genomic dimerization and potentially contribute to subsequent homologous recombination events. However, further studies are required to explore such functions.

Our study subsequently investigated the proposed hijacking of cellular miRNA-1307-3p by the s2m element of SARS-CoV-2 (47-49). Here, we have shown for the first time experimentally that the SARS-CoV-2 s2m binds multiple copies of miRNA-1307-3p, and furthermore, that this interaction is affected by the state of s2m dimerization. The role of host cellular miRNA hijacking by viruses has been demonstrated extensively in previous studies, typically benefitting the viral life cycle. Bovine viral diarrhea virus, for instance, interacts with miRNA-17 upon infection, resulting in enhanced replication, translation, and genomic RNA stability (70). Similarly, miRNA-133a is bound by the 3’ UTR of the Dengue virus, essentially “sponging” it from its normal function of regulating the polypyrimidine tract-binding (PTB) protein (71). This interaction results in an upregulation of the PTB protein, leading to increased viral replication and translation. Here, through analogy to these other systems, we suggest a similar mechanism with SARS-CoV-2 hijacking cellular miRNA-1307-3p through the s2m element. Further experiments are required to test this mechanism, but if proven correct, this would result in upregulated expression levels of its target mRNAs, including those that encode for various interleukins (IL18) and interleukin receptors (IL6R, IL10RA, IL10RB) linked to the “cytokine storm” involved in ARDS development in severe COVID-19 patients (25, 50, 51). Interestingly, IL6, IL10, and IL18 are secreted by ATII epithelial cells, the primary targets for SARS-CoV-2 infection (53). The latter of these cytokines, IL18, has further been suggested to be at the “eye of the cytokine storm”, acting synergistically with various other primary cytokines in a proinflammatory manner to modulate the expression of secondary cytokines, such as IFNγ (72–74). Moreover, overproduction of IL18 has been linked to a predisposition for macrophage activation syndrome, resulting in an overactive immune response, as well as Kawasaki disease and systemic lupus erythematosus, autoimmune diseases which present symptoms similar to those seen in critically ill paediatric COVID-19 patients (75, 76). In addition to its role of promoting increased inflammatory response, high levels of IL18 have been also shown to be correlated with COPD, hypertension, cardiovascular disease, and gestational diabetes mellitus, all of which enhance patient risk of experiencing severe COVID-19 symptoms (75, 77). Thus, we contemplate the implications of a fine-tuned molecular switch from s2m element dimerization to miRNA-1307-3p binding, with the latter mechanism potentially aiding in severe COVID-19 symptom development. Altogether, the results of this study implicate dimerization of the highly conserved s2m element of SARS-CoV-2 as a potentially significant molecular target for antiviral therapeutics for COVID-19 as well as future coronavirus-related disease outbreaks.

## SUPPLEMENTARY DATA

Supplementary Data are available at NAR online.

## NIST DISCLAIMER

Certain commercial equipment, instruments, and materials are identified in this paper in order to specify the experimental procedure. Such identification does not imply recommendation or endorsement by the National Institute of Standards and Technology, nor does it imply that the material or equipment identified is necessarily the best available for the purpose.

