## Supplemental figures for "SARS-CoV-2 highly conserved s2m element dimerizes via a kissing complex and interacts with host miRNA-1307-3p"

### Supplemental Figure S1

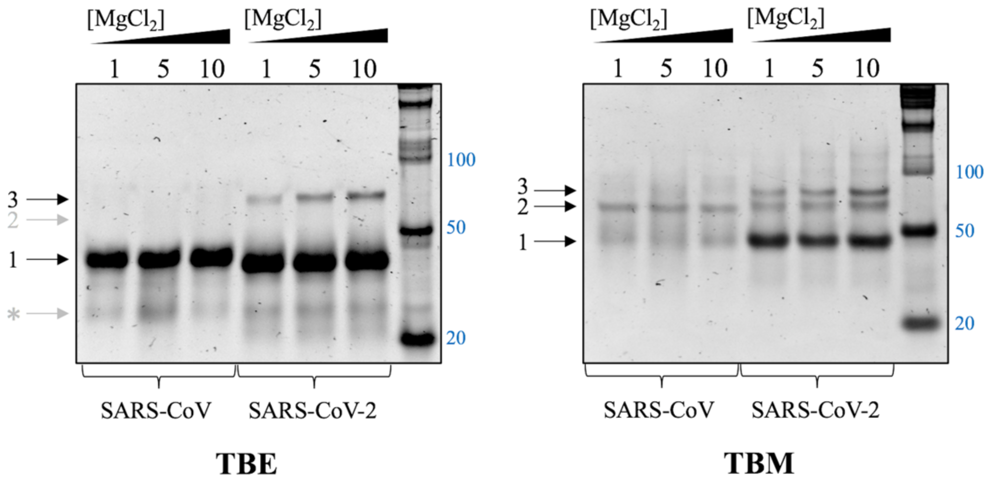


**Supplemental Figure S1.** Nondenaturing TBE and TBM PAGE of SARS-CoV and SARS-CoV-2 following 24-hour incubation. Samples were incubated in the presence of 1, 5, and 10 mM $MgCl_2$ followed by incubation for 24-hours at room temperature. SARS-CoV-2 s2m undergoes spontaneous conversion from the kissing dimer conformation (arrow 2) to a thermodynamically stable extended duplex (arrow 3), which is present even in the TBE gel (left panel), while SARS-CoV s2m is unable to undergo this spontaneous conversion. Both SARS-CoV and SARS-CoV-2 undergo degradation during 24-hour incubation, as denoted by an asterisk (*). A low molecular weight RNA ladder (Abnova) was used as a reference.

### Supplemental Figure S2

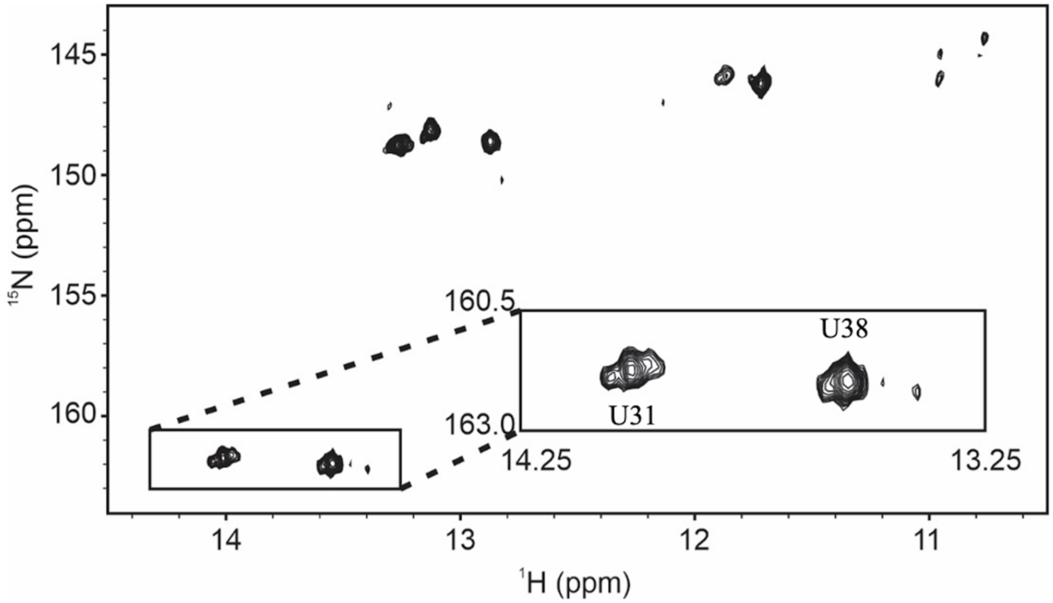


**Supplemental Figure S2.** Imino region of the $^{1}H$,$^{15}N$ SOFAST-HMQC of the wild-type SARS-CoV-2 s2m construct. The spectrum was collected at $^{15}N$ natural isotopic abundance and was recorded at 900 MHz at 25 °C. Inset: Expansion of the Uracil region showing the multiple conformers of U31.

### Supplemental Figure S3

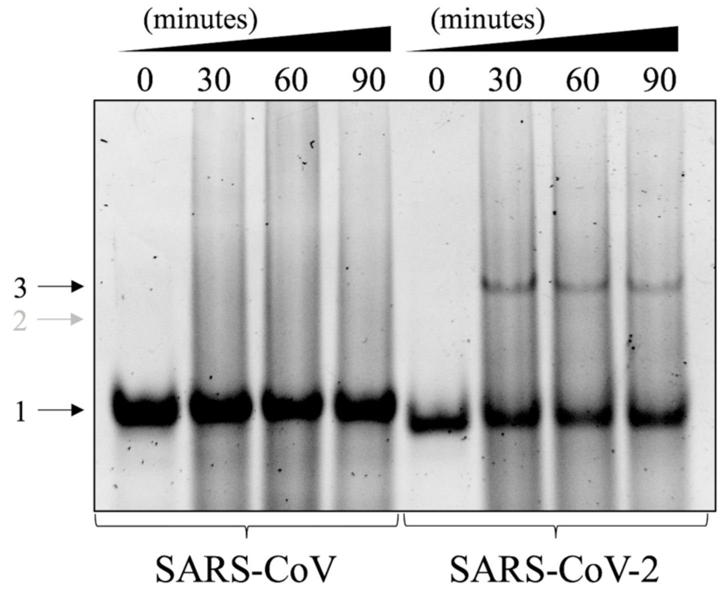


**Supplemental Figure S3.** Time-dependent incubation of s2m elements with the SARS-CoV-2 N protein. Both SARS-CoV and SARS-CoV-2 s2m elements were incubated in the presence of 1 mM $MgCl_2$ and the SARS-CoV-2 N protein at a 1:2 ratio over the course of 90-minutes, followed by electrophoresing on a nondenaturing TBE gel. While SARS-CoV-2 s2m is converted to a thermodynamically stable extended duplex conformation (arrow 3), SARS-CoV exists solely in its monomeric conformation (arrow 1).

**Supplemental Figure S4**

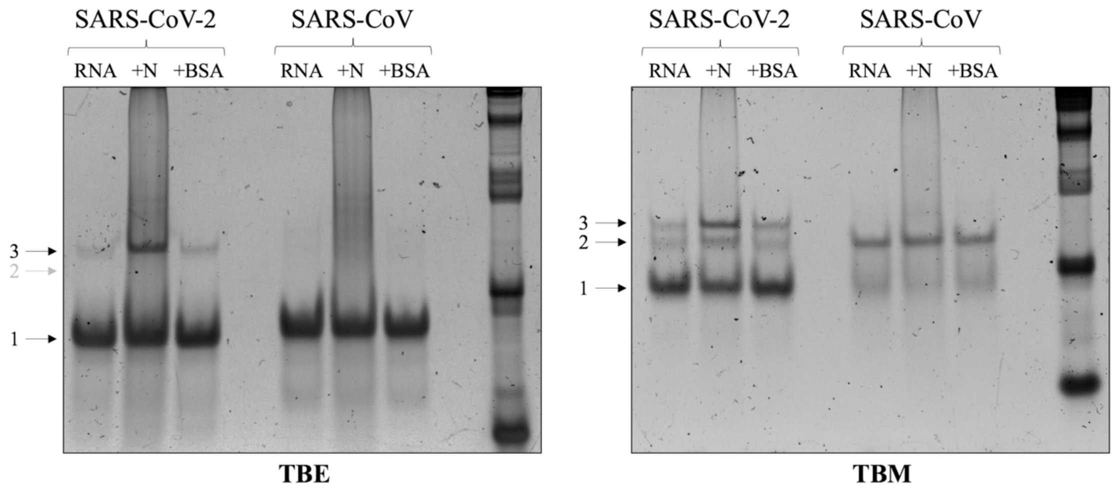


**Supplemental Figure S4.** Chaperone activity of viral N protein at biologically relevant temperature. Both SARS-CoV and SARS-CoV-2 s2m elements were incubated in the presence of 1 mM $MgCl_2$ for 1-hour, followed by addition of the N protein in a 1:2 ratio and additional incubation for 30-minutes at 37 °C. Proteinase K was added to digest the N protein prior to samples being split and electrophoresed by TBE and TBM nondenaturing PAGE. The kissing complex formed by SARS-CoV-2 s2m (arrow 2) is converted to a thermodynamically stable extended duplex (arrow 3) by chaperone activity of the N protein and accelerated by higher temperature compared to Figure 6, whereas SARS-CoV s2m is unable to undergo similar conversion. Control samples in which both s2m elements were incubated in the presence of BSA in a 1:2 ratio revealed no changes in conversion to stable duplex conformation.

**Supplemental Figure S5**

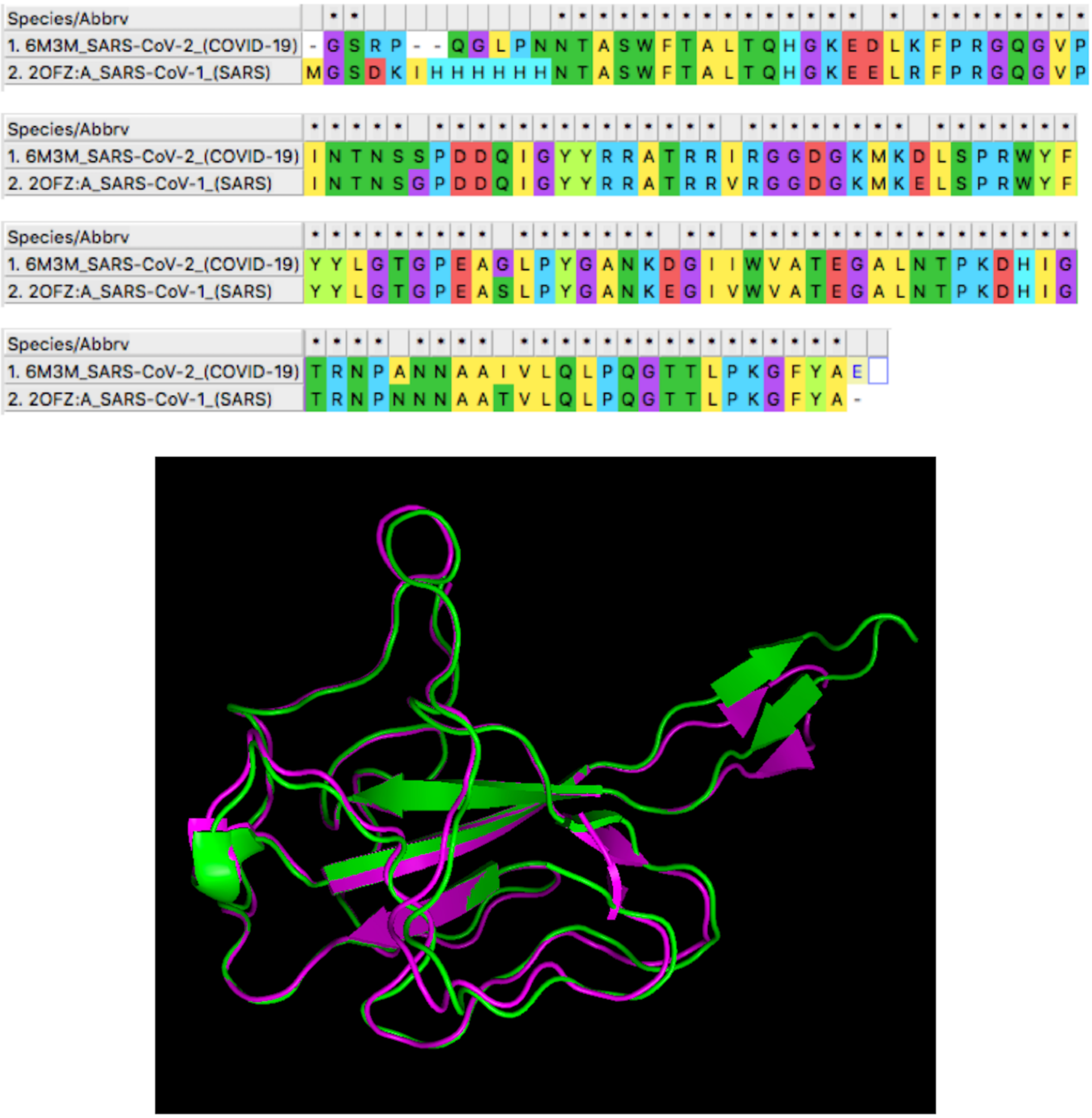


**Supplemental Figure S5.** Sequence and structure similarity between SARS-CoV and SARS-CoV-2 N protein RNA-binding domains (RBDs). Sequences were compared using MEGA X molecular alignment software (top). Structural alignments of SARS-CoV N protein RBD (PDB: 1SSK; green) and SARS-CoV-2 N protein RBD (PDB: 6M3M; purple) were performed using PyMol visualization software (bottom).

**Supplemental Figure S6**

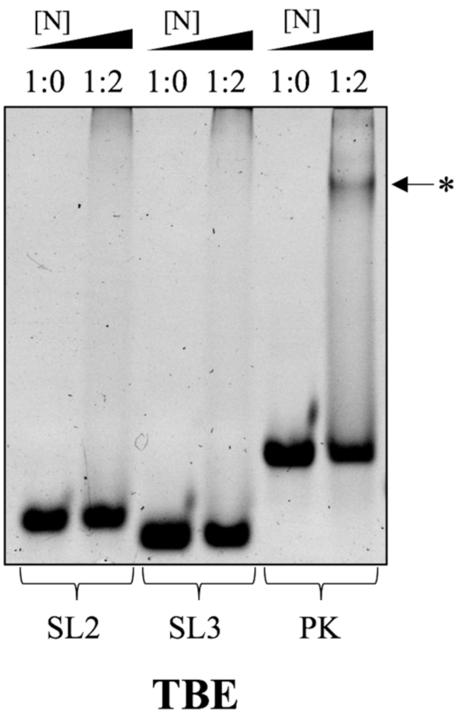


**Supplemental Figure S6.** SARS-CoV-2 N protein binding to stem-loop controls. SL2, SL3, and PK stem-loop elements of the SARS-CoV-2 genome were incubated in the presence of 1 mM $MgCl_2$ and the SARS-CoV-2 N protein in 1:0 and 1:2 ratios. Samples were electrophoresed by nondenaturing TBE PAGE, revealing the N protein does not bind to SL2 or SL3, and weakly binds to the PK element, with the faint complex band denoted by an asterisk (*).

**Supplemental Figure S7**

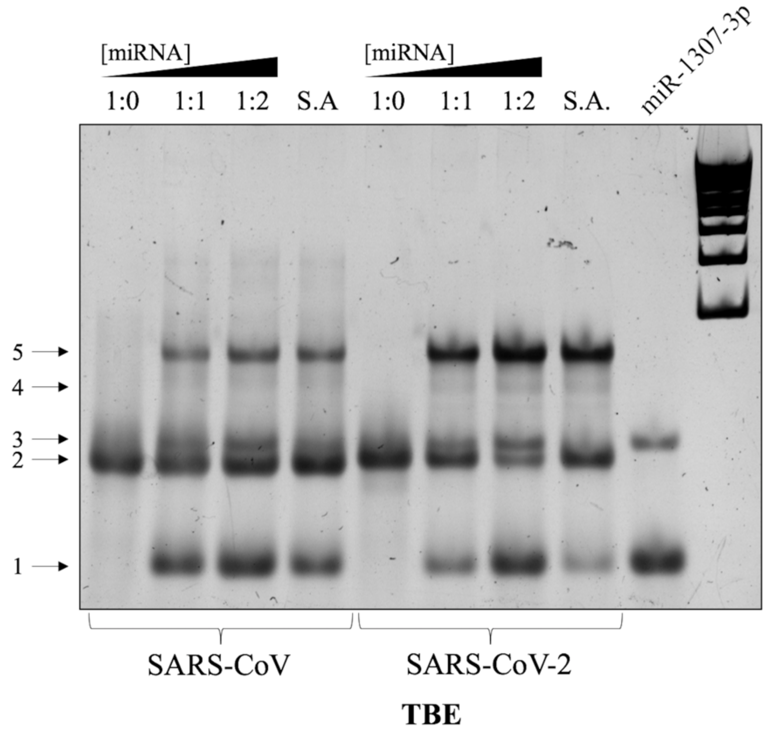


**Supplemental Figure S7.** Interaction of the SARS-CoV and SARS-CoV-2 s2m elements with host cellular miRNA-1307-3p. Both s2m elements at concentrations of 1 µM were incubated with 1 mM $MgCl_2$ and miRNA-1307-3p at increasing ratios (up to 1:2) for 1-hour. Samples were analyzed by TBE nondenaturing PAGE. Upon chelation of $Mg^{2+}$ ions, both SARS-CoV and SARS-CoV-2 bind to one (arrow 4) and two (arrow 5) molecules of miRNA-1307-3p, with an increased intensity seen for the SARS-CoV-2 complex bands compared to SARS-CoV and a concomitant decrease in monomeric SARS-CoV-2 s2m (arrow 2). Control lanes of each s2m element slow-annealed (S.A.) with miRNA-1307-3p at a 1:2 ratio did not indicate greater binding, but instead increased conversion of monomeric miRNA-1307-3p (arrow 1) to its dimeric conformation (arrow 3).
